# Dynamic inositol pyrophosphate synthesis is a targetable therapeutic opportunity in ovarian cancer

**DOI:** 10.64898/2026.08.25.747159

**Authors:** Daniel P Bondeson, David Husselbee, Saoirse Hanbury, Amirshayan Chadeganipour, Gabriel Mesa, Alison Cameron, Janhavi Y Sawant, Carly Langan, Tuhina Bhattacharya, YuhJong Liu, Hajer Siala, Erin M Swanson, Mustafa Kocak, Karthik Srinivasan, Blanche C Ip, Nancy Dumont, Randall Burton, Jennifer A. Roth, David E Root, John G Doench, Alexandra E Gould, Dean Proctor, Todd R Golub

**Affiliations:** UMass Chan Medical School, Department of Systems Biology, Worcester MA, USA; The Broad Institute, Cambridge MA, USA; Department of Pediatric Oncology, Dana-Farber Cancer Institute, Harvard Medical School, Boston, MA, USA

## Abstract

We previously reported that the phosphate exporter XPR1 is required to prevent toxic phosphate accumulation in ovarian cancer cells. To guide therapeutic development, we sought to systematically compare potential strategies to inhibit XPR1: directly targeting the phosphate efflux channel, targeting its partner protein KIDINS220, or inhibiting the synthesis of inositol pyrophosphates (PP-InsPs), metabolites which activate XPR1. We evaluated functional domains in XPR1 and KIDINS220 using mutational scanning and found that loss of function mutations in XPR1 clustered in distinct regions throughout the protein, with the most deleterious mutations in the PP-InsP-binding domain. In contrast, loss of function mutations in KIDINS220 were infrequent and altered the localization of XPR1, consistent with a scaffolding role for KIDINS220. These data highlight the functional relevance of PP-InsPs, which we confirmed by inhibiting their synthesis using IP6K inhibitors. We demonstrate that IP6K inhibition phenocopies XPR1 inhibition across hundreds of cancer cell lines, with the mechanism of sensitivity solely due to inhibition of cellular phosphate efflux. Finally, we show that IP6K inhibitors decrease tumor burden in xenograft models of ovarian cancer, but that the rapid resynthesis of PP-InsPs requires high exposures to achieve efficacy. This study comprehensively evaluates the XPR1-dependent phosphate efflux network and reinforces the concept of directly targeting XPR1 as a precision medicine strategy to benefit patients with ovarian cancer.

## INTRODUCTION

Survival rates for ovarian cancer have not significantly improved in the past twenty years: new insights into its etiology and novel therapeutic strategies are needed. Strategies that complement current standard of care and have non-overlapping mechanisms of resistance would be particularly beneficial. We recently demonstrated that ovarian cancers frequently exhibit altered phosphate homeostasis due to overexpression of the phosphate importer SLC34A2/NaPi-2b^1,2^. Although much therapeutic effort has been focused on developing antibody-drug conjugates targeting SLC34A2^3,4^, our recent work demonstrated that the excessive phosphate import caused by SLC34A2 creates a synthetic reliance on cellular phosphate export. By inhibiting the phosphate export complex consisting of XPR1 and KIDINS220, we showed that ovarian cancer cells accumulate toxic amounts of inorganic phosphate, suggesting that targeting cellular phosphate homeostasis could be a selective therapeutic opportunity in these cancers. Nevertheless, much work is needed to identify the ideal therapeutic approach to take advantage of altered phosphate homeostasis in ovarian cancer.

Although much work has focused on understanding organismal phosphate homeostasis in mammals, cellular homeostasis is less characterized. Cellular uptake is mediated by the ubiquitously expressed importers PiT1/PiT2 -- which also have roles in sensing extracellular phosphate -- and tissue-specific importers in the SLC34 family^5^. Phosphate abundance inside the cell is primarily sensed metabolically through the synthesis of inositol pyrophosphates^6^, although other metabolic sensors also exist in specific tissues^7^. The intracellular signaling networks affected by inorganic phosphate and inositol pyrophosphates are also poorly characterized^8^, but it has been shown that inositol pyrophosphates regulate the activation of XPR1-dependent phosphate efflux^9–12^. Recent work has added structural details to the inositol pyrophosphate-mediated regulation of the XPR1:KIDINS220 phosphate efflux complex^13–15^. In addition, the precise contribution of KIDINS220 to phosphate homeostasis is poorly understood. It was originally identified as a scaffolding protein for receptor tyrosine kinase signaling, and loss of function mutations cause a rare neurodevelopmental syndrome^16^ (SINO). The contribution of phosphate efflux to this syndrome is unknown. In addition, our data suggest a positive role for KIDINS220 in maintaining proper surface localization of XPR1^1^, whereas recent reports suggest that KIDINS220 stabilizes the inactive conformation of XPR1^13^.

From this brief survey, much remains to be understood about cellular phosphate homeostasis, implying that there could be several different mechanisms to target phosphate homeostasis in ovarian cancer. Here, we sought to systematically compare potential therapeutic modalities to target cellular phosphate homeostasis, including disruption of the XPR1:KIDINS220 protein complex and depletion of PP-InsPs. Using mutational scanning, we identify the SPX domain of XPR1 as the most functionally intolerant component of the phosphate efflux complex. In addition, using chemical probes which deplete PP-InsP, we demonstrate that inhibition of IP6K and XPR1 both achieve selective anti-cancer activity, with shared mechanisms of sensitivity but distinct mechanisms of resistance.

## RESULTS

### Mutational scanning of the XPR1:KIDINS220 protein complex

We first sought to systematically identify regions of the XPR1:KIDINS220 protein complex that are critical for ovarian cancer fitness. XPR1 is 696 amino acids and consists of a phosphate channel (also termed an EXS domain), a transmembrane ‘core’ responsible for structural stability, and a gating SPX domain^13,15^. KIDINS220 is much larger – 1,771 amino acids – and consists of an N-terminal ankyrin-repeat domain, transmembrane domains which include a potential Walker P NTPase domain, and a C-terminal disordered domain (Figure 1A, top). Deletion of the C-terminal domain of KIDINS220 has been shown to cause a neurodevelopmental syndrome, and the Ankyrin repeat has been shown to bind to XPR1 and stabilize its inactive conformation ^13^, but the contributions of these diverse functions have not been studied in the context of ovarian cancer survival. To compare the functional consequence of each domain, we used CRISPR-based mutational scanning to evaluate the functional consequences of many mutations in parallel^17–19^. We used ABE8e, a dCas9-fusion capable of installing adenosine-to-guanosine mutations (A.G)^18^ to introduce mutations in endogenous *XPR1* and *KIDINS220*; loss of function mutations should manifest as loss of fitness in ovarian cancer cell lines and subsequent depletion of the sgRNA from the population. We designed a library of sgRNA targeting every possible NG PAM of each gene: 1,011 sgRNA targeting XPR1 and 2,435 sgRNA targeting KIDINS220. It is not possible to perfectly predict editing outcomes for Abe8e, but we predict this library has the potential for editing ∼75% of amino acids with >2,700 total edits (6% of all possible amino edits, Extended Data Figure 1A, B). Because OVISE cells are equally dependent on both *XPR1* and *KIDINS220*^1^, the fitness defect of each sgRNA can be compared between and within genes to identify functionally relevant domains. We scored each sgRNA by comparing its relative distribution within the population of cells at day 21 versus its initial representation at day 4; more depletion suggests more functional relevance (Figure 1A-B and Extended Figure 1C-E).

**Figure 1:**
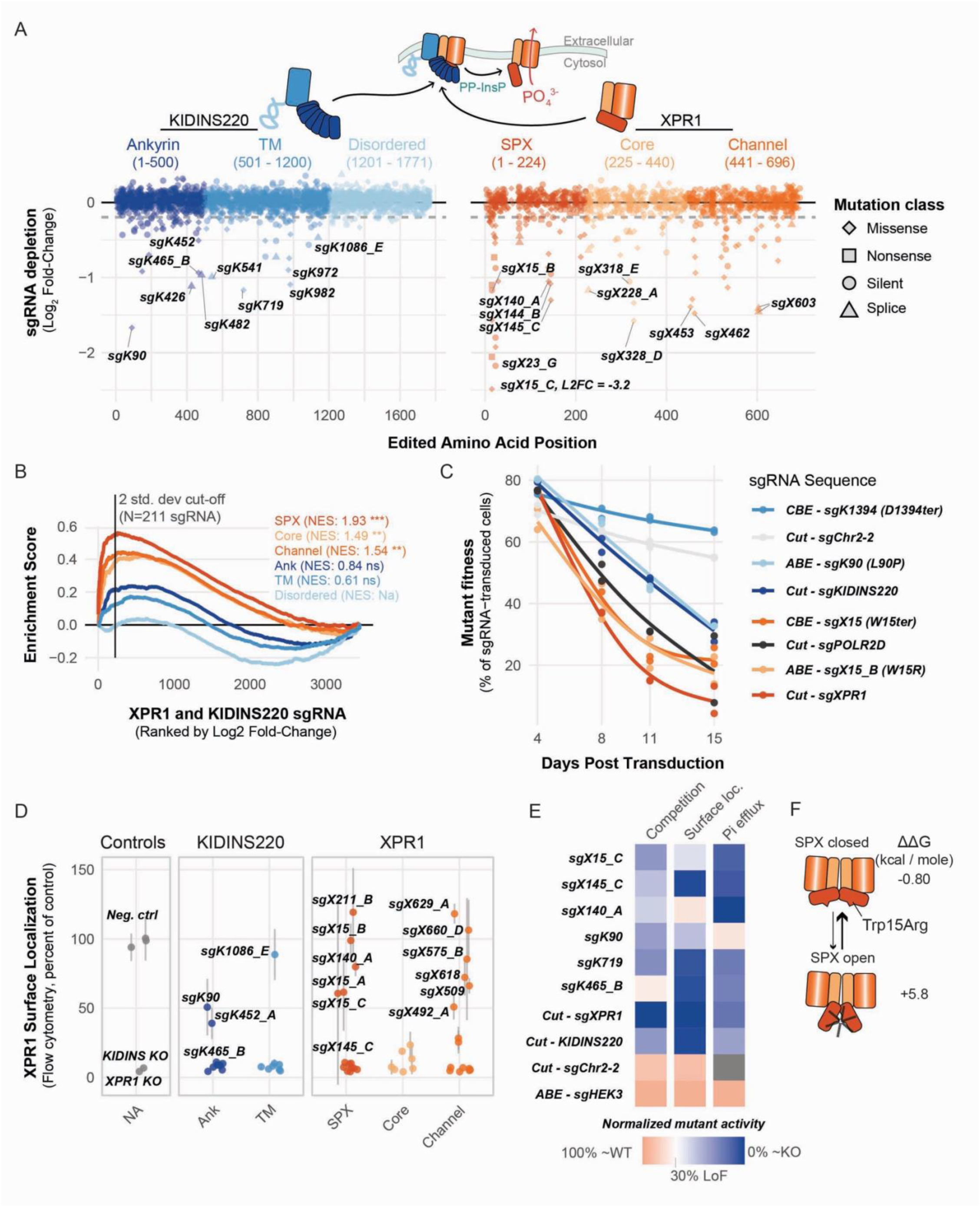
Mutational scanning of the phosphate efflux protein complex prioritizes the regulatory SPX domain of XPR1. A, Top, schematic of the XPR1:KIDINS220 phosphate efflux complex. Bottom, results of mutational scanning of the protein complex. A library of sgRNA was introduced into OVISE ovarian cancer cells along with the Abe8e A.G editor; each sgRNA is named for the target (X for XPR1, K for KIDINS220), the predicted amino acid edited, and a unique ID if multiple sgRNA target the same site. The cells were then cultured for two weeks, and sgRNA abundance was assessed at day 4 and day 21. Shown is the depletion of each sgRNA, averaged across three replicates. The dotted line represents two standard deviations away from the median depletion of negative control sgRNA. Separately, an APOBEC-based C.T editor was introduced with the same sgRNA library and only results for sgRNA predicted to introduce nonsense mutations are displayed. Note that sgX15 scored with a value of −3.2 and is displayed higher for visual clarity. B, Domain-level enrichment analysis for sgRNA across XPR1 and KIDINS220. sgRNA targeting the indicated domains are ranked by depletion score (X axis), and the relative enrichment of each domain compared with a random distribution is shown on the Y axis. Shown in parenthesis are the normalized enrichment scores and significance as assessed by a permutation test (*** p < 0.001, ** p < 0.01, ns not significant). C, Validation of top scoring base editing sgRNA in competition experiments. sgRNA and Cas enzymes were introduced into OVISE-eGFP cells and then mixed with OVISE-mCherry cells on day 4; the eGFP/mCherry ratio was determined on the day of mixing and on the indicated days. The cells were then cultured for 2 weeks, and the GFP/mCherry ratio was measured at each split. CBE, cytosine base editor BE3.9 with sgRNA designed to install nonsense mutations; ABE, adenine base editor Abe8e with predicted edits indicated; Cut, wildtype Cas9 enzyme to introduce double stranded breaks in the indicated genes. sgChr2-2 targets a gene desert on Chromosome 2. D, Analysis of scoring sgRNA in HEK293T cells. Cell surface expression of XPR1 was assessed seven days after transfection of each sgRNA and Cas enzyme into HEK293T cells. The mean fluorescence intensity from two separate transfections is plotted with error bars representing the standard error. E, In addition to the competition (C) and surface localization (D) experiments, cellular efflux of 32P-labeled inorganic phosphate was evaluated in HEK293T cells. Each assay is summarized in the heatmap with orange values indicating wildtype-levels of activity and blue values indicating the activity of XPR1-inactivated cells; white values indicate 30% loss in protein function. ABE – sgHEK3 installs an A.G mutation at a gene desert. F, The highest scoring Trp15Arg mutation is predicted to selectively destabilize the open conformation of XPR1, leading to an auto-inhibited phosphate channel regardless of cellular InsP8 levels. Trp15Arg stability was calculated with FoldX in comparison to PDBs 9JXG (closed) and 9JXH (open).

At the gene-level, we found that the most depleted sgRNAs targeted XPR1 rather than KIDINS220: 74 significantly depleted sgRNA target XPR1 compared to 53 for KIDINS220, despite only 29% of the initial library being designed to target XPR1 (Figure 1A, Extended Data Figure 1E). We next calculated domain-level enrichment scores, asking whether domains were enriched for highly depleted sgRNA (Figure 1B). We found that the most mutationally intolerant domain was the N-terminal SPX domain of XPR1, followed by the other domains of XPR1; strongly depleted sgRNA were not enriched across KIDINS220 domains.

To understand how individual mutations cause loss-of-viability, we validated >50 individual sgRNAs that were significantly depleted. We tested each sgRNA in competition-based fitness assays in the OVISE cell line, and then further studied cellular localization, protein stability, and phosphate transport assays using transient transfection in HEK293T cells which do not require XPR1 for survival (Figure 1C-E). There was a high degree of concordance between the primary screen and validation in the competition assay (Figure 1C). Consistent with prior efforts^20,21^, most loss of function mutations either destabilize or prevent proper trafficking of the protein complex, as measured by decreased cell surface staining with the viral XPR1-binding protein X-RBD. Nearly all KIDINS220 mutations decreased cell surface localization, further suggesting its role as a scaffold protein without relevant enzymatic activities. In contrast, many of the SPX- and channel-targeting sgRNA were properly localized (Figure 1D).

Of the individual sgRNA tested, few mutations had no effect on protein levels, and we validated that these mutations decrease cellular phosphate efflux activity (Figure 1E). These include sgX140, predicted to introduce the same Leu140Pro mutation observed in patients with primary brain calcification due to XPR1 loss of function^22^. In addition, the most potently depleted sgRNA in the library did not destabilize XPR1 protein levels and was predicted to introduce a Trp15Arg mutation that has not been reported before (Figure 1E,F and Extended Data Figure 1F). This mutation resides in the SPX domain which undergoes large structural rearrangements upon binding to inositol pyrophosphates and opening of the phosphate permeation channel^13^. We noticed that Trp15 was oriented towards solvent in the closed conformation (apoXPR1-SPX3, PDB 9JXG) while in the open conformation (PP-InsP8-XPR1-SPX2, PDB 9JXH), Trp15 was buried and made several important contacts to stabilize the phosphate channel gate. To validate this, we performed *in silico* prediction of the stability of the Trp15Arg mutant and found larger protein destabilization for the open versus the closed conformation (ΔΔG of +5.8 kcal/mol versus −0.80 kcal/mol).

The relatively few scoring mutations targeting *KIDINS220* was surprising given the relevance of *KIDINS220* in a variety of neurodevelopmental syndromes. For example, the truncating mutation Gln1393ter has been shown to hinder neuronal development^23^, but this mutation was not depleted in a second base editor screen using a dCas9-BE3.9 editor capable of introducing C.T mutations (Extended Data Figure 1D), nor in validation competition assays. This mutation had no effect on XPR1 cell surface localization or cellular phosphate transport. Together, these observations suggest that *KIDINS220* is pleiotropic and that its role in phosphate transport is not responsible for *KIDINS220*-liniked disorders.

### IP6K1/2 inhibitors show SLC34A2^HIGH^-selective anti-cancer activity

Our mutagenesis experiment demonstrated that the regulatory SPX domain was as functionally intolerant to mutations as the phosphate permeation channel itself and complete genetic inactivation of *XPR1*. The SPX domain is an auto-inhibitory gate which keeps the phosphate channel closed^9,11,12^; upon binding to inositol pyrophosphates (PP-InsPs, most notably 5-diphosphoinositol pentakisphosphate, or 5-PP-InsP_7_, and 1,5-bisdiphosphoinositol tetrakisphosphate, or 1,5-PP-InsP_8_) the SPX domain is stabilized, resulting in an open phosphate channel^13,14^. Inositol pyrophosphates are in turn synthesized in response to rising cellular phosphate levels: As cellular phosphate levels rise, ATP synthesis is increased and inositol pyrophosphate-synthesizing enzymes can utilize the higher concentrations of ATP. The IP6K1 and IP6K2 kinases phosphorylated InsP_6_ to form the pyro-phosphorylated molecule 5-PP-InsP_7_ which is further phosphorylated to yield 1,5-PP-InsP_8_ by the PPIP5K1 and PPIP5K2 kinases (Figure 2A). Thus, the inositol pyrophosphate metabolic network connects cellular inorganic phosphate concentrations to activation of XPR1.

**Figure 2:**
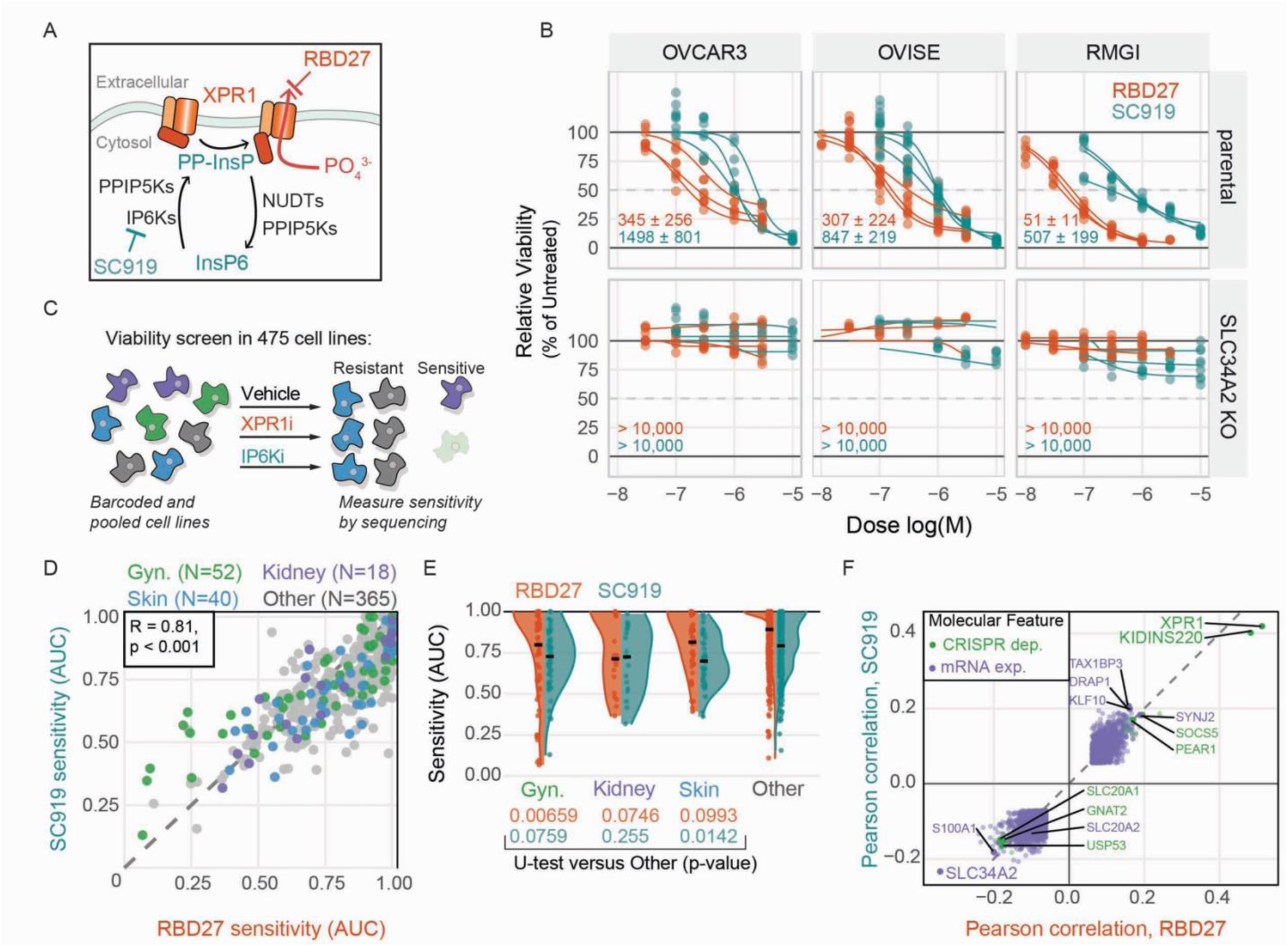
XPR1 and IP6K activities are functionally redundant across 100s of cancer cell lines. A, Cartoon representation of the inositol pyrophosphate pathway, which activates XPR1 in response to elevated cellular phosphate levels. RBD27 (orange) is a viral protein-based inhibitor of XPR1, while SC919 (teal) is a small molecule inhibitor of IP6K1, IP6K2, and IP6K3. B, Viability profiling of XPR1 (RBD27, orange) and IP6K (SC919, teal) inhibitors. Three isogenic pairs of ovarian cancer cell lines (parental or *SLC34A2*-inactivated) were treated with the indicated doses of drug for 7 days with redosing on day 4. Viability was assessed via CellTiterGlo, and the viability relative to untreated control wells is plotted on the Y-axis. Each curve represents a non-linear regression fit; three representative replicates are displayed. The inset shows the calculated IC50 as the mean of the displayed replicates +/- standard deviation. C, Schematic of chemogenetic profiling for XPR1 and IP6K inhibitors using the PRISM assay. D, Comparison of RBD27 and SC919 in the PRISM assay against 475 cancer cell lines. Each compound was tested in 6 doses, and the area under the dose response curve (AUC) is plotted for each cell line. E, Summary of sensitivity data from D comparing gynecological cancer (Gyn., ovarian and uterine, N = 52), kidney cancer (N=18), skin cancers (N=40) compared to all other cancer types. Below, results from a Mann-Whitney U test are displayed for RBD27 (orange) and SC919 (teal). F, RBD27 and SC919 sensitivity profiles (AUC profiles) were correlated with gene expression (mRNA expression, purple) or gene essentiality (CRISPR dependency Chronos scores, green) profiles for each cell line. mRNA expression and gene essentiality profiles were obtained from DepMap Public Release 25Q3.

To test whether PP-InsP depletion is toxic to SLC34A2-expressing ovarian cancer cell lines, we used SC919, a recently published small molecule dual inhibitor of IP6K1 and IP6K2^24^. To compare the activity of IP6K inhibition with direct XPR1 inhibition, we also used a RBD27, a proof-of-concept XPR1 inhibitor based on the viral receptor binding domain of the Xenotropic Murine Leukemia Virus which uses XPR1 for viral entry^25,26^ (Figure 2A, Extended Data Figure 2A-E). As we have shown previously^1^, XPR1 inhibition completely suppresses the viability of *SLC34A2*-expressing, but not isogenic *SLC34A2*-inactivated cell lines (Figure 2B). Satisfyingly, SC919 demonstrates similar potency and selectivity (Figure 2B) in the same panel of cell lines and could induce cell death in an SLC34A2-dependent manner (Extended Data Figure 3A).

We next asked whether IP6K inhibition phenocopied XPR1 inhibition or if PP-InsPs have other cellular roles beyond activating XPR1. Although >1,000 cancer cell lines have been profiled for their sensitivity to *XPR1* inactivation, sensitivity to inactivation of *IP6K1* and *IP6K2* is likely masked by paralog buffering: accordingly, both genes are not essential in any cancer cell lines profiled in DepMap (Extended Data Figure 3B). Instead, we used the PRISM assay^27,28^: a highly multiplexed cancer viability assay in which 475 cancer cell lines were genetically barcoded, pooled together, and then exposed to a dose series of RBD27 and SC919 (Figure 2C-F, Extended Data Figures 3C-F). Consistent with a shared mechanism of action, the drugs were highly correlated in their anti-cancer activity (R = 0.81, p<0.001, Figure 2D) but uncorrelated to other drugs profiled in the PRISM assay^28^ (Extended Data Figure 3F), suggesting a unique pattern of anti-cancer activity. Ovarian and uterine cancer lineages were significantly more sensitive to RBD27 and SC919 than other lineages, along with slight enrichment for kidney and skin cancer (Figure 2E). Gene essentiality for *XPR1* and *KIDINS220,* and high expression of *SLC34A2,* were the most correlated molecular features across all cell lines that are sensitive to RBD27 or SC919 (Figure 2F and Extended Data Figure 3D, E). Therefore, inositol pyrophosphates have no functional role in maintaining cancer cell fitness aside from activating XPR1-dependent cellular phosphate efflux.

### IP6K inhibitors decrease tumor burden *in vivo*

We next sought to evaluate the in vivo efficacy of targeting cellular phosphate homeostasis. The RBD27 protein was cleared rapidly from mice after tail vein injection (Extended Data Figure 4A), precluding further in vivo study. In contrast, oral administration of SC919 demonstrated a pharmacokinetic profile with good plasma exposures and a long-half life when formulated with 5% DMSO in 20% hydroxy-propyl-beta-cyclodextrin (HPβCD formulation, Figure 3A and Extended Data Figure 4B). Doses of SC919 that should achieve tumor drug coverage above the viability IC90 were well tolerated in mice, with only minor weight loss observed (Figure 3B).

**Figure 3:**
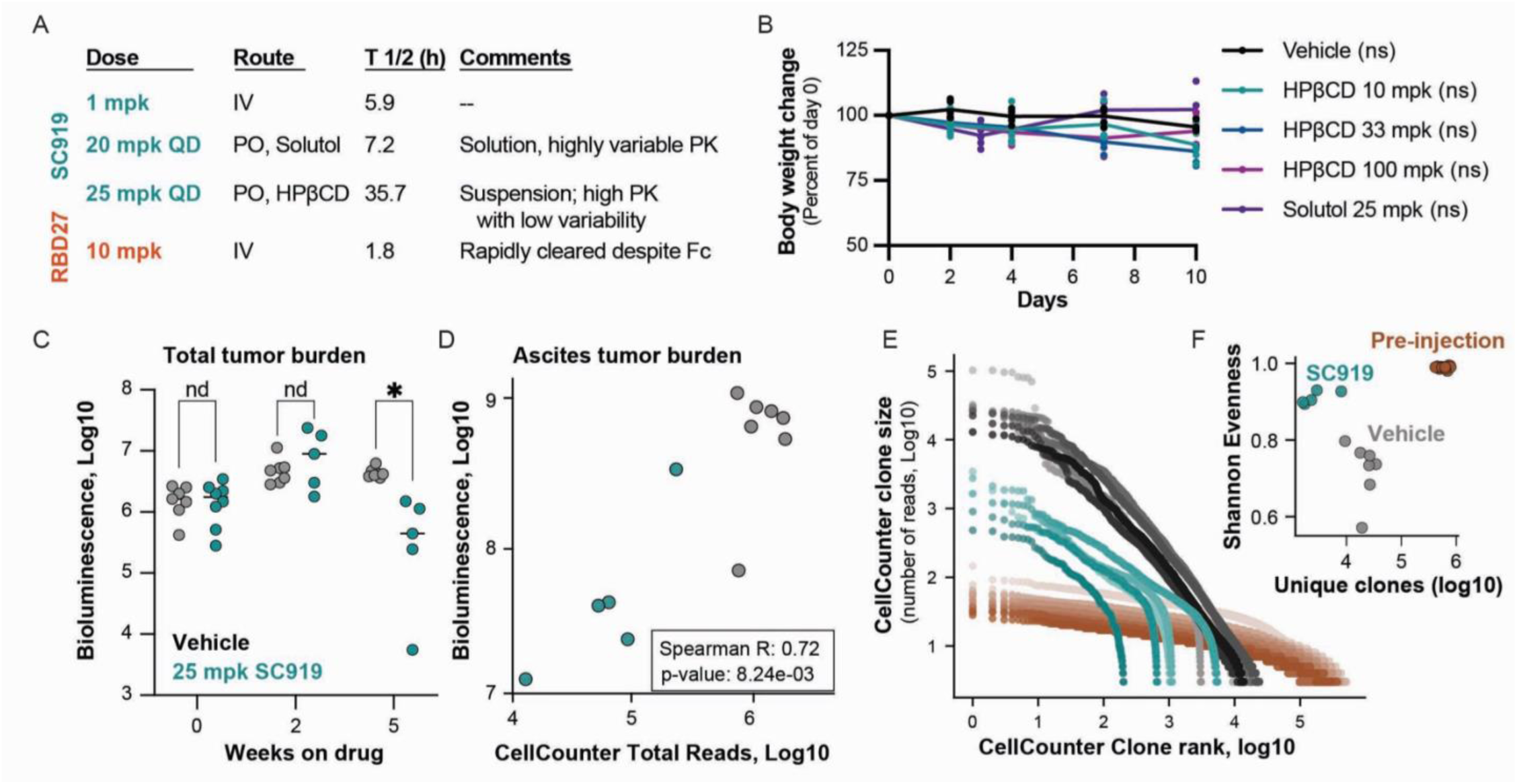
IP6K inhibition decreases tumor burden in vivo. A, Dosing regimen and pharmacokinetic half-life of SC919 after tail vein injection (IV), oral gavage (PO) delivery in the Solutol formulation, or oral gavage delivery of the hydroxy-prolyl-beta-cyclodextrin (HPβCD) formulation. B, The HPβCD formulation permits dosing at high MTD doses, as measured by animal body weight over ten days of dosing. ns, not significant as assessed by one-way ANOVA comparing the day 10 body weights between vehicle and treatment groups C, OVISE cells expressing luciferase and CellCounter Barcodes were injected into the peritoneal cavity of NSG mice. After 10 days of tumor growth, the animals were treated daily with vehicle or 25 mg/kg SC919 in the cyclodextrin formulation and bioluminescence was assessed throughout the study. * p<0.01 by ANOVA. D, At the end of the OVISE study, animals were dissected and tumor tissues were examined ex vivo for bioluminescence (Y-axis). In parallel, genomic DNA was isolated and CellCounter barcodes were quantified (X-Axis). E, Clonality analysis of tumor cells from the OVISE study. At the end of the study, genomic DNA from the Ascites was isolated and CellCounter clonality barcodes were analyzed by next generation sequencing. Each point represents a unique barcode, with the size of that clone (number of reads) represented on the Y-axis and the rank of that clone for all clones within an individual mouse on the X-axis. F, Quantification of population size (total number of CellCounter clones) and diversity (Shannon evenness, where a value of 1 is a perfectly even population).

To test the anti-cancer efficacy of SC919, we developed a model of disseminated ovarian carcinomatosis. OVISE cells were engineered to express luciferase and injected into the intraperitoneal cavity of immunocompromised mice and allowed to grow for one week before the animals were treated with SC919 (Extended Data Figure 3C-D). After several weeks of dosing, we observed significantly decreased whole-body bioluminescence in SC919-treated animals (Figure 3C, Extended Data Figure 4D). Upon termination of the study and ex vivo analysis of different organs, we observed that many tumor cells were still present in the intraperitoneal fluid (i.e. ascites), and that this site had the largest drug-induced decrease in bioluminescent signal (Figure 3D, Extended Data Figure 4E).

Dosing regimen that inconsistently achieve coverage above theIC90 may lead to selection for and outgrowth of intrinsically resistant clones. To assess this, we had also introduced a high complexity library of genetic barcodes to identify and track clones into the OVISE cells shortly before injection in mice (Figure 3D-F, Extended Data Figure 4C, F). We refer to this system as CellCounter, as it offers an orthogonal metric to understand *in vivo* tumor evolution and clonal dynamics (Extended Data Figure 4F). We estimated that animals injected with 6,000,000 cells received approximately 600,000 unique molecular identifiers (UMIs), a marker of clonality (Figure 3E-F). At the end of the study, we extracted genomic DNA from ascites fluid and organ sites, and quantified UMI diversity using next generation sequencing. We found that the total read counts across all UMIs within an individual tumor was highly correlated to the bioluminescence of that organ (Figure 3D), demonstrating the robustness of our sequencing protocol. We found that drug-treated animals had 10X fewer unique CellCounter clones (average of ∼12,000 in the vehicle-treated animals compared to ∼1,200 in the SC919-treated animals, Figure 3E-F). We also observed an increase in the evenness of the distribution of CellCounter barcode in drug-treated tumors compared to vehicle-treated tumors, suggesting that drug treatment prevents outgrowth of specific clones (Figure 3E, F). In addition, we also noted decreased dissemination of clones from the ascites to other organ sites (Extended Data Figure 4F). Together, these data suggest that drug treatment both decreased the proliferation of rapidly expanding clones – increasing evenness – and caused cell death in some clones – decreasing the number of unique clones.

### IP6K inhibition requires high exposure to achieve anti-tumor efficacy

To further explore potential resistance mechanisms to XPR1 or IP6K inhibition, we conducted genome-scale modifier screens (Figure 4A and Extended Data Figure 5A and B). After lentiviral transduction of an enCas12a genome-scale sgRNA library^29^, cells were cultured for two weeks with RBD27 or SC919 at doses which reduced proliferation to a similar extent (Extended Data Figure 5A). We sequenced the sgRNA and compared the relative abundance of each sgRNA in the drug-treated populations to the untreated population: enrichment indicates drug resistance and depletion indicates drug sensitization. The drugs displayed highly correlated chemogenetic profiles (Figure 4A, Extended Data Figure 4B), with *SLC34A2* inactivation conferring strong resistance (Figure 4A). Interestingly, sgRNA targeting *XPR1* were strongly enriched in RBD27-treated cells, potentially due to outgrowth of XPR1 mutants that are resistant to RBD27-induced inhibition. Despite their overall similar activity, we noted that inactivation of *NUDT4* was unique in its enrichment in the SC919-treated condition but not the RBD27-treated cells (Figure 4A, B). *NUDT4* encodes a diphosphoinositol polyphosphate phosphohydrolase (DIPP2) enzyme that is capable of dephosphorylating inositol pyrophosphate and thus opposes the activity of IP6K1/2^30^. This result suggests that the synthesis and degradation of PP-InsPs is highly dynamic in ovarian cancer cells, consistent with recent reports^31^. To test this, we evaluated cellular phosphate efflux in a washout experiment: although washout of RBD27 from cells displayed sustained efflux inhibition, washout of SC919 from cells almost immediately restored cellular phosphate efflux (Figure 4C). Therefore, IP6K inhibition is more easily reversed from cells compared to XPR1 inhibition.

**Figure 4:**
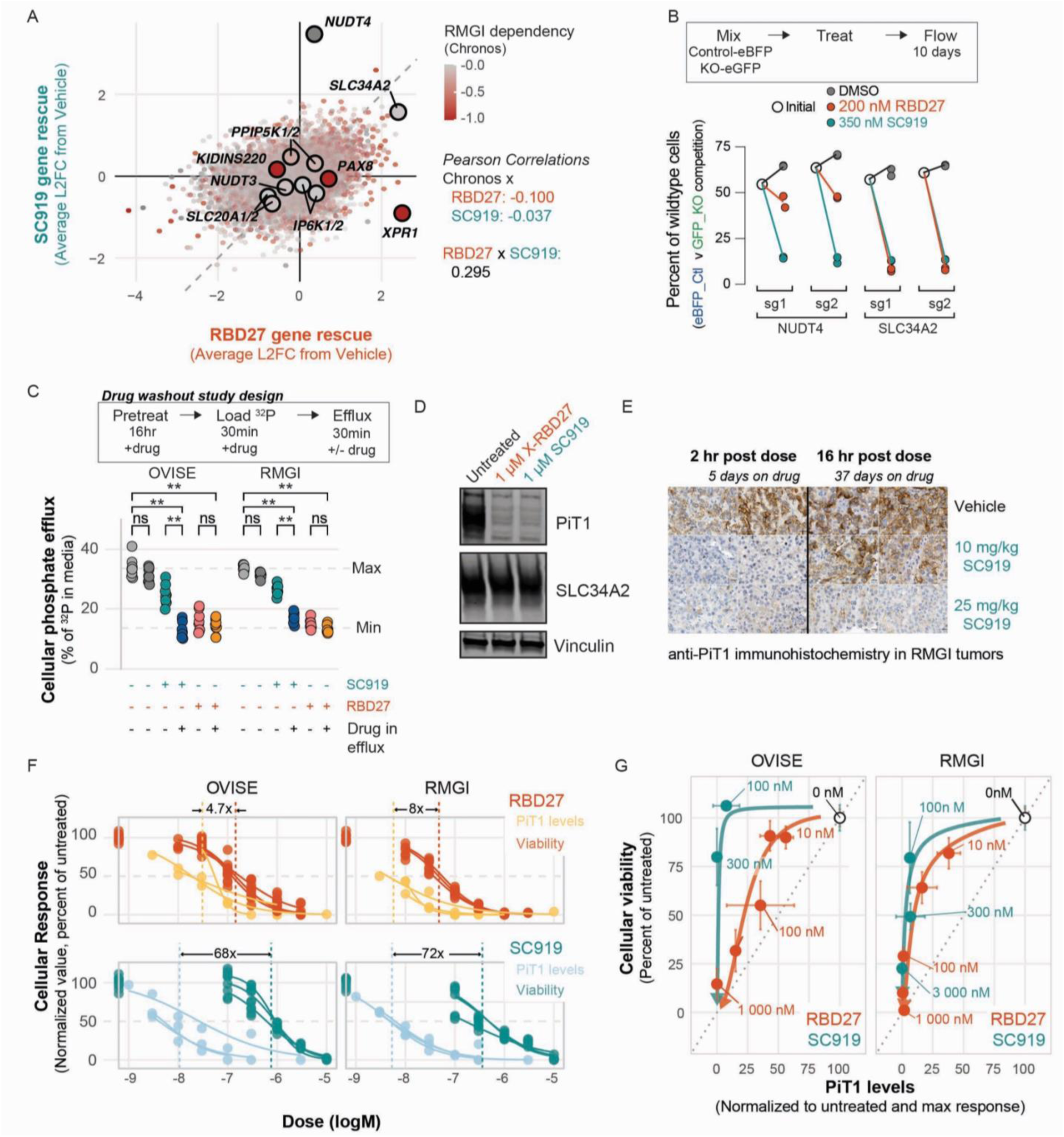
IP6K inhibitors require high exposures for sustained inhibition of XPR1. A, Genome-scale modifier screen for RBD27 and SC919. The Humagne Set C and Set D genome-scale CRISPR/enCas12a libraries were introduced via lentiviral transduction into RMGI cells, and then the cells were cultured in the presence of a vehicle control, 250 nM RBD27, or 350 nM SC919 for 14 days. The Log2Fold-Change enrichment of each gene drug-treated conditions relative to vehicle is plotted on the X-Axis for RBD27 and the Y-Axis for SC919. Each point is colored by the gene essentiality (Chronos score) for that gene. Larger points indicate known regulators of phosphate homeostasis and inositol pyrophosphate metabolism. Right, Pearson correlation between each value is indicated. B, Validation of the inositol pyrophosphate polyphosphatase NUDT4 as a unique mechanism of resistance to IP6K inhibition. GFP-labeled gene knockouts (NUDT4 or SLC34A2) were mixed with BFP-labeled control cells and cultured with the indicated concentrations of drugs. C, IP6K inhibitors require constant exposure to inhibit cellular phosphate efflux. After pre-treatment with RBD27 or SC919 and a 30-minute pulse with ^32^P-labeled inorganic phosphate, cells were washed and incubated in drug-containing or drug-replete media to initiate phosphate efflux. After 25 minutes, ^32^P activity was measured in the conditioned media and the lysate, and phosphate efflux is calculated as the percentage of ^32^P in the conditioned media. Significance was assessed by two-way ANOVA. **p<0.001, ns not significantD, Suppression of the PiT1 phosphate importer after 24h treatment with XPR1 and IP6K inhibitors in the RMGI ovarian cancer cell line. E, Solid tumor tissues from the RMGI xenografts in C were evaluated for PiT1 expression levels using immunohistochemistry. N=2 mice per dosing group. Note that the 2-hour post dose animals were part of the “PD group” which was only dosed for 5 days, whereas the 16 hour post dose animals were part of the “survival group” which had received SC919 for 5 weeks. F, Comparison of the potency of SC919 and RBD27 to suppress PiT1 after 24 hours and cellular viability after 7 days of drug exposure (re-dosing on day 4). All datasets were scaled between 100% for untreated response (which are plotted at the far left of the plot) and 0% for the maximal, saturating response observed. Each curve represents an independent experiment. Dotted lines represent the calculated IC50, and the difference between PiT1 and viability IC50s is noted above each graph. G, Pharmacodynamic comparison of SC919 and RBD27. The modeled dose-response curves from F were used to simulate the relationship between PiT1 suppression and cellular viability, which is displayed as the trajectory according to increasing dose to the lower left. Data points represent shared doses between both assays, displayed as the mean +/- the standard deviation of at least three replicates.

Our observations that IP6K inhibition is rapidly reversible suggest that consistently high drug exposure, over the IC90 threshold, is required to inhibit tumor growth. To test this, we evaluated the efficacy of a second formulation of SC919 that demonstrated a shorter half-life (Solutol, Figure 3A and Extended Data Figure 4B) and is predicted to only intermittently expose tumors to >IC90 concentrations upon daily dosing. We observed no efficacy in reducing RMGI tumor growth (Extended Data Figure 5C, D), despite equivalent sensitivity of OVISE in vitro (Figure 2). To evaluate tumor exposure, we developed a pharmacodynamic assay based on suppression of the phosphate importer PiT1^1,10^ (Figure 4D). RMGI tumor tissues demonstrated robust PiT1 suppression 2 hours after SC919 dosing that returned to untreated levels within 16 hours (Figure 4E). To better understand this result, we compared the in vitro potency of PiT1 degradation at 24 hours and relative cellular viability after 7 days (Figure 4F, G). We noted that IP6K inhibition via SC919 demonstrated a 72-fold difference between PiT1 degradation and reduced cellular viability. In contrast, XPR1 inhibition via RBD27 demonstrated only an 8-fold difference between PiT1 degradation and reduced viability. Together, these data suggest that the rapid resynthesis of inositol pyrophosphates requires constant inhibition of IP6K enzymes to achieve the same level of anti-cancer activity as partial but direct inhibition of XPR1.

## DISCUSSION

In this work, we demonstrate the importance of carefully and comprehensively surveying the complex network of enzymes and proteins that are required for essential cellular processes. Specifically, we compared XPR1, KIDINS220, or IP6K enzymes as ideal therapeutic targets in SLC34A2^HIGH^ cancers. Using mutational scanning, we highlight the functional relevance of XPR1 and the scaffolding roles for KIDINS220: a much lower rate of scoring sgRNA in KIDINS220, the lack of clustering of those sgRNA, and the observation that the most profound sgRNA decrease cell surface expression of XPR1 all point to a localization/scaffold function for KIDINS220. In addition, the observation that phosphate efflux is unaffected by mutations in KIDINS220 that cause neurodevelopmental disorders suggests pleiotropic roles for KIDINS220. More work is needed on the physiological functions of XPR1, KIDINS220 and the function of their protein complex across diverse tissues.

Using chemical probes and extensive pharmacologic profiling, we demonstrate that XPR1 and IP6K are nearly functional equivalent in terms of cancer survival. The poly-pharmacology of SC919 (i.e. inhibiting both IP6K1 and IP6K2) demonstrates remarkably specific anti-cancer activity, as evidenced by selective toxicity across hundreds of cell lines and a shared chemogenetic profile with the XPR1 inhibitor RBD (Figures 2 and 4). Moreover, IP6K inhibition decreased tumor burden in SLC34A2^HIGH^ tumors at tolerated doses, suggesting there is a therapeutic window between efficacy and on-mechanism toxicity. The specific reliance of ovarian cancers on IP6K activity suggest that SLC34A2^HIGH^ tumors have increased PP-InsP levels to support cellular phosphate flux via XPR1. Nevertheless, the rescue of SC919 by *NUDT4* inactivation suggests that PP-InsP levels are highly dynamic, as reported recently^31^. Future work should focus on profiling the highly interconnected and redundant phosphate homeostasis pathways and their dysregulation in cancer.

Although this work highlighted similarities between IP6K and XPR1 inhibition, an important question remains as to whether an improved IP6K inhibitor (or inhibitors for other members of the PP-InsP pathway like PPIP5Ks) would yield a more favorable therapeutic window than direct inhibitors of XPR1. Importantly, the highly dynamic synthesis and degradation of PP-InsPs likely enables normal cells to rapidly respond to fluctuations in phosphate availability and/or metabolism but requires high drug exposures to achieve marginal anti-tumor efficacy (Figure 3C-F). Such high dosing may manifest in undesirable off-target toxicities due to unselective inhibitors. In addition, PP-InsPs likely have additional functions in other tissues aside from XPR1 inhibition. For example, recent studies in mice have highlighted unique functions for Ip6k1/2 and Xpr1 in maintaining normal kidney function^32,33^. Taken together, direct inhibition of XPR1 is likely a favorable therapeutic strategy relative to IP6K inhibition, although further in vivo studies using optimized XPR1 inhibitors will be needed to robustly address this important question. Future work should focus on deepening our understanding of XPR1:KIDINS220, and phosphate homeostasis more generally, to fully realize the promise of precision cancer medicine.

---

**Extended Data Figure 1:**
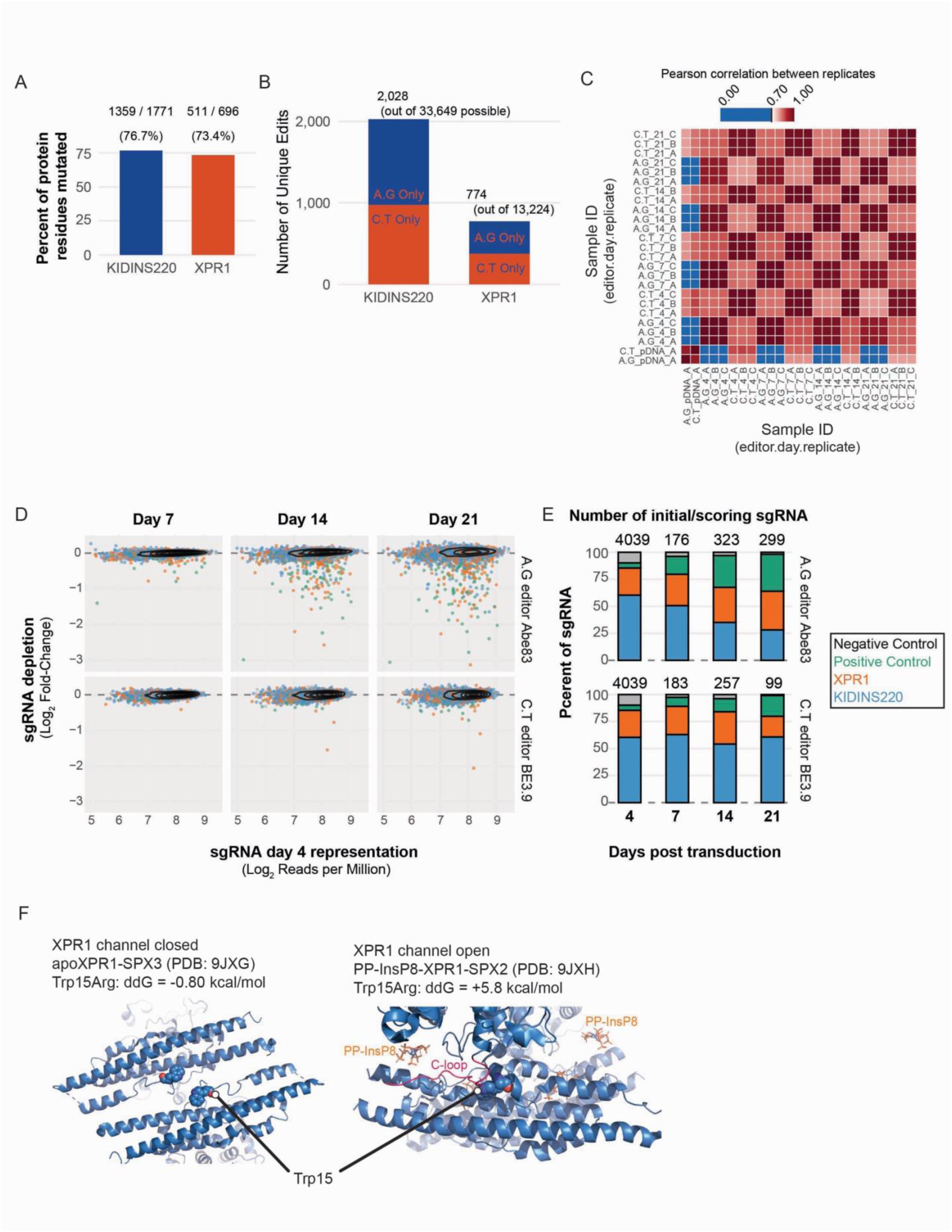
Mutational scanning of the XPR1:KIDINS220 protein complex in ovarian cancer cells. A, Residue coverage of the base editor library. B, Number of unique edits introduced by both the A.G and C.T editors, compared with the theoretical ‘saturating’ library containing all possible variants. C, Pearson correlation of the normalized reads per million between individual replicates samples in the base editing CRISPR screen for XPR1:KIDINS220. D, Depletion of sgRNA over time (Log_2_FC from day 4 to day 21) versus initial representation (Log_2_RPM) highlighting that depletion was not skewed by initial library representation. Note that the C.T editor had much lower depletion and sgRNA scoring rates. E, Representation of scoring sgRNA. The day 4 stacked bar graph represents the initial library of 4,039 sgRNA; days 7, 14, and 21 are the proportion of scoring sgRNA (>2 standard deviations from the median of the negative control sgRNA). Note the enrichment of positive control and XPR1 sgRNA in the A.G but not the C.T library. F, Zoom-in visualizations of the atomic structure of the SPX domain of XPR1 as determined by Wang et al., 2025. Trp15Arg stability was calculated with FoldX in comparison to PDBs 9JXG (closed) and 9JXH (open).

**Extended Data Figure 2:**
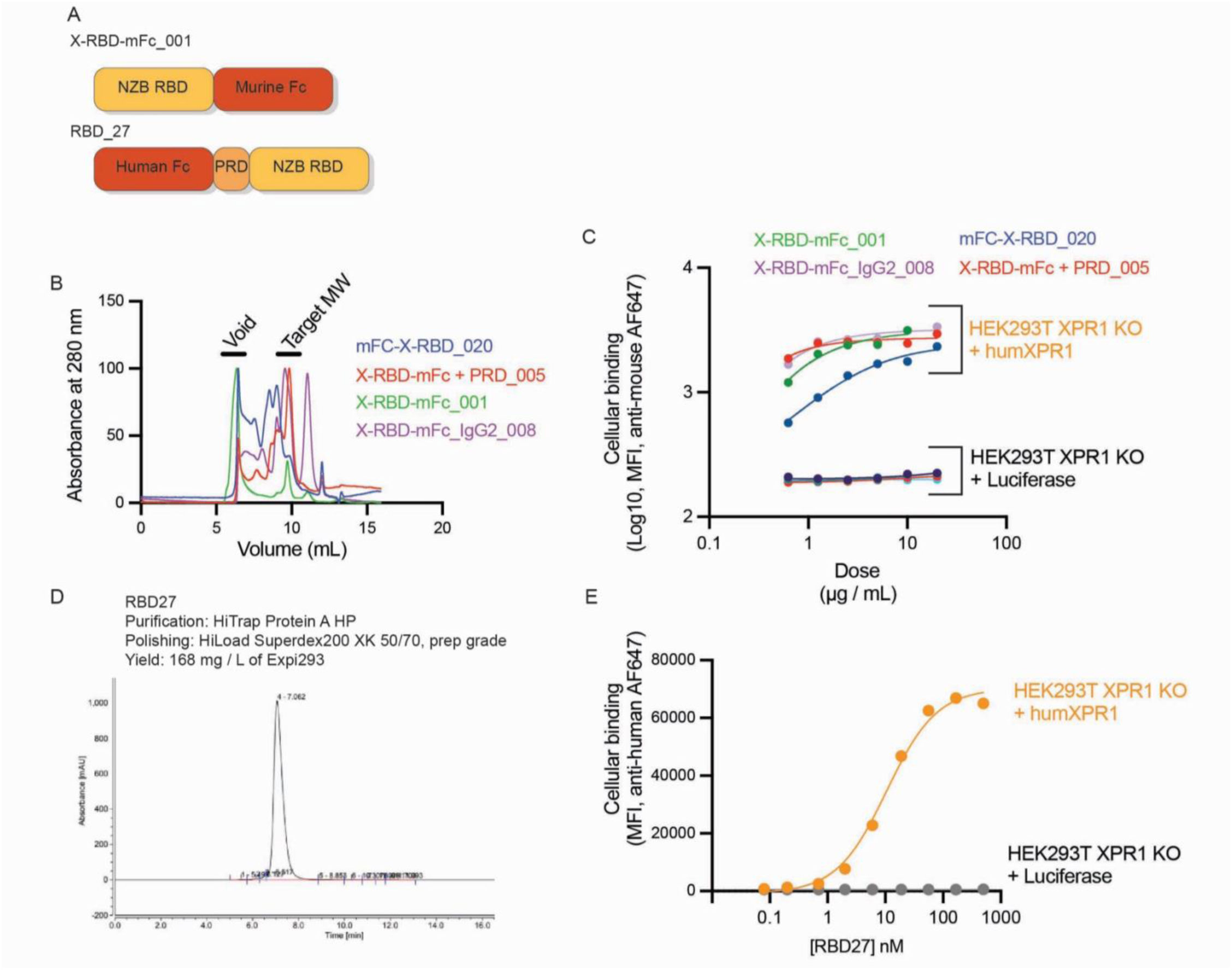
Development of RBD27 for cellular pharmacology. A, Schematic illustration of the original X-RBD protein compared with RBD27. The Receptor Binding Domain (RBD) protein is derived from the NZB murine leukemia virus and in initial publications was purified with a C-terminal murine IgG1 Fc tag. Instead of a C-terminal murine Fc tag, RBD27 has an N-terminal human Fc tag and contains the proline-rich domain (PRD). B, Size exclusion chromatography of X-RBD fusion proteins after protein A purification. Protein 001 represents the original construct (panel A), Protein 005 contains the proline-rich domain as a linker between the RBD and Fc tag, Protein 008 contains a murine Fc tag derived from IgG2, and Protein 020 is a N-terminal Fc fusion of X-RBD. C, Binding of X-RBD fusion proteins to HEK293T (expressing endogenous XPR1) or BaF3 cells. Note that it has been previously shown that X-RBD fusion proteins do not bind to murine XPR1. D, Size exclusion chromatography of RBD27 after purification with Protein A and polishing with size exclusion chromatography. E, Binding of RBD27 to HEK293T cells expressing human XPR1 compared to XPR1 knockout cells. RBD27 binds with comparable affinity as X-RBD_001 but with a much cleaner SEC profile (Panel D).

**Extended Data Figure 3:**
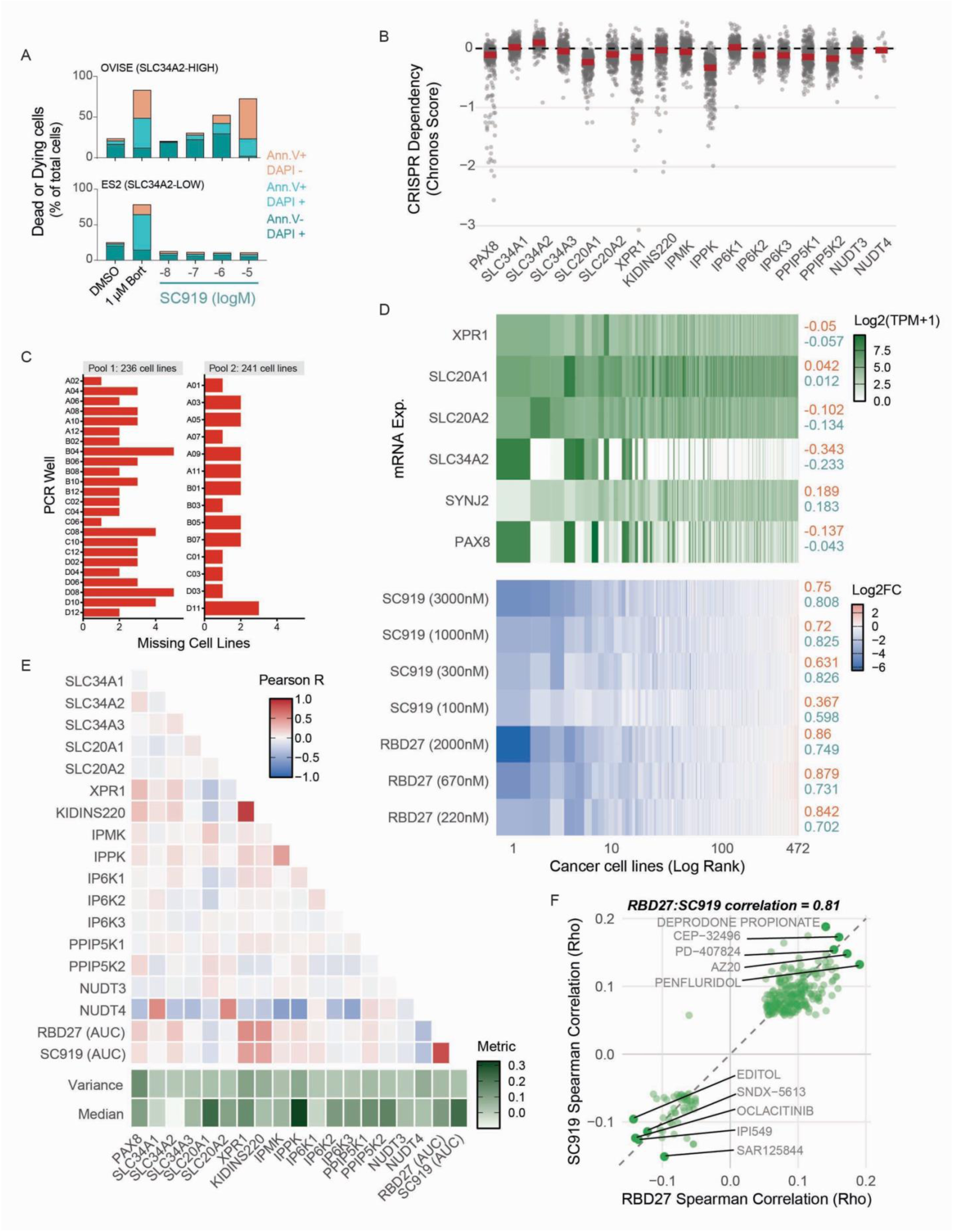
Inhibition of IP6K1 and IP6K2 selectively induces cell death in SLC34A2-HIGH and XPR1-dependent cancer cell lines. A, SLC34A2-specific cell death with SC919. After 6 days of treatment with SC919 or 24 hours of Bortezomib (Bort) at the indicated doses, OVISE and ES2 cells were collected and stained with Annexin V (to detect apoptotic cells) and DAPI (to detect dead, permeabilized cells) followed by analysis by flow cytometry. B, Paralogs likely buffer IP6K and PPIP5K enzymes from essentiality across cell lines. Displayed are the Chronos dependency scores for each gene across the DepMap Public 25Q3 dataset (N = 1,100 cell lines except for NUDT4). C, Cell line recovery from the PRISM experiment. After 6 days of treatment with RBD27 or SC919 (see Figure 2), cell line genomic barcodes were amplified and sequenced. The entire collection of cells is divided into 12 sub-pools which were collapsed into two pools each containing ∼240 cell lines for PCR and sequencing. 5 or fewer cell lines were missing from each PCR reaction, indicating good recovery. D, Correlation of gene expression and drug sensitivity. The top correlated mRNA expression profiles are plotted for the 475 cancer cell lines in the PRISM assay. Cell lines are ranked by average sensitivity to RBD27 and SC919 and plotted on a log-rank scale to display the most sensitive cell lines. To the right of each heatmap, Pearson correlation coefficients with RBD27 (orange) and SC919 (teal) sensitivity are indicated. E, Correlation of gene dependency and drug sensitivity. The correlation of each gene dependency profile (panel B) and drug sensitivity is displayed. Below, the variance and median of each killing profile is displayed, calculated on the absolute value of the Chronos score such that larger positive median values indicate stronger fitness defects observed in more cell lines. Note that correlations to *NUDT4* dependency are likely spurious due to the small number of cell lines profiled for this gene. Strongest correlations are observed for *XPR1*, *KIDINS220*, RBD27, and SC919, but not genes encoding inositol pyrophosphate metabolic enzymes. F, Correlation of RBD27 and SC919 with other drugs profiled in PRISM as part of drug repurposing studies.

**Extended Data Figure 4:**
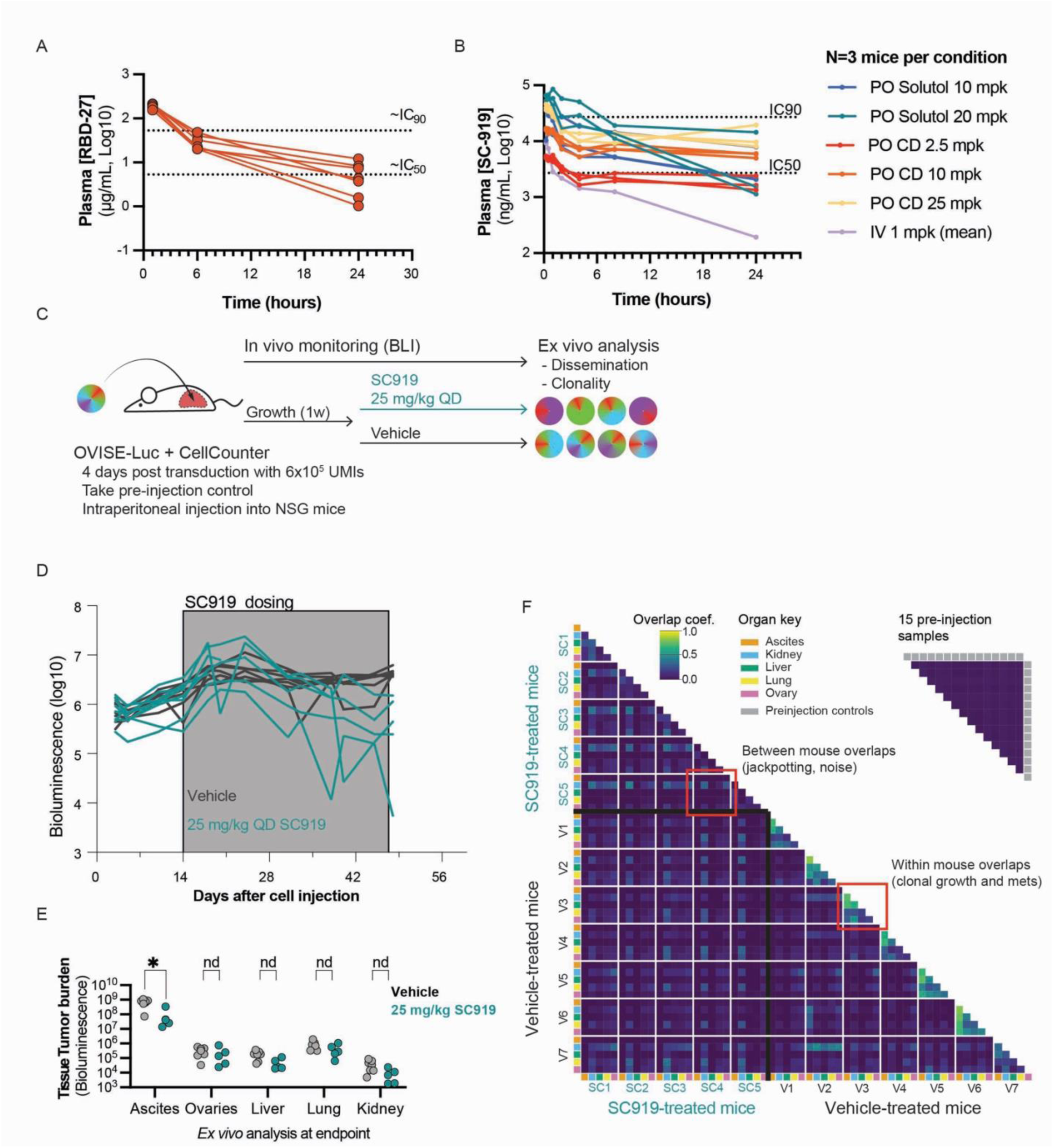
IP6K inhibition decreases tumor burden in vivo. A, Pharmacokinetic profiles of RBD27 after tail vein injection. 6 mice were injected with RBD27, and plasma was isolated at 15 minutes, 6 hours, and 24 hours later. RBD27 was measured in cells via ELISA directed to the human IgG1 Fc tag. IC50 and IC90 values were calculated from Figure 2B and are not corrected for bioavailability. B, Pharmacokinetic profiles of SC919 after tail vein injection (IV), oral gavage (PO) delivery in the Solutol formulation, or oral gavage delivery of the hydroxy-propyl-beta-cyclodextrin (HPβCD) formulation. IC50 and IC90 values were calculated from Figure 2B and are corrected for plasma protein binding (99.1% in RPMI + 10% FBS and 99.96% in plasma) C, Study schematic to assess IP6K inhibitor efficacy in a model of disseminated ovarian carcinomatosis. OVISE cells were engineered to express luciferase for bioluminescent tracking, as well as a CellCounter barcoding library to evaluate clonal evolution. These cells were injected into the peritoneal cavity of NSG mice and bioluminescence was measured twice per week. After 2 weeks of tumor growth, the animals were dosed via oral gavage daily with 25 mg/kg of SC919 formulated in 5% DMSO in 20% HPβCD in water. At the end of five weeks of dosing, mice were sacrificed and disseminated tumor burden was assessed using bioluminescence and next generation sequencing. D, Spaghetti plot of whole-body tumor burden in individual mice. E, Ex vivo analysis of bioluminescence across diverse organs. At the end of the study, each animal was sacrificed the indicated organs were isolated, washed twice with PBS, and then bioluminescence was measured. Note that almost all tumor burden was observed in the ascites fraction. F, CellCounter barcodes enable tracking the evolution of tumor clonality across different animals. After sacrifice, dissection, and ex vivo bioluminescent imaging (panel E), CellCounter barcodes were quantified with next generation sequencing. The heatmap shows the overlap coefficient (how many CellCounter barcodes are in both populations relative to the overall population size) for each sample. Note that inter-mouse overlap is generally very low relative to intra-mouse overlap in the Vehicle-treated animals.

**Extended Data Figure 5:**
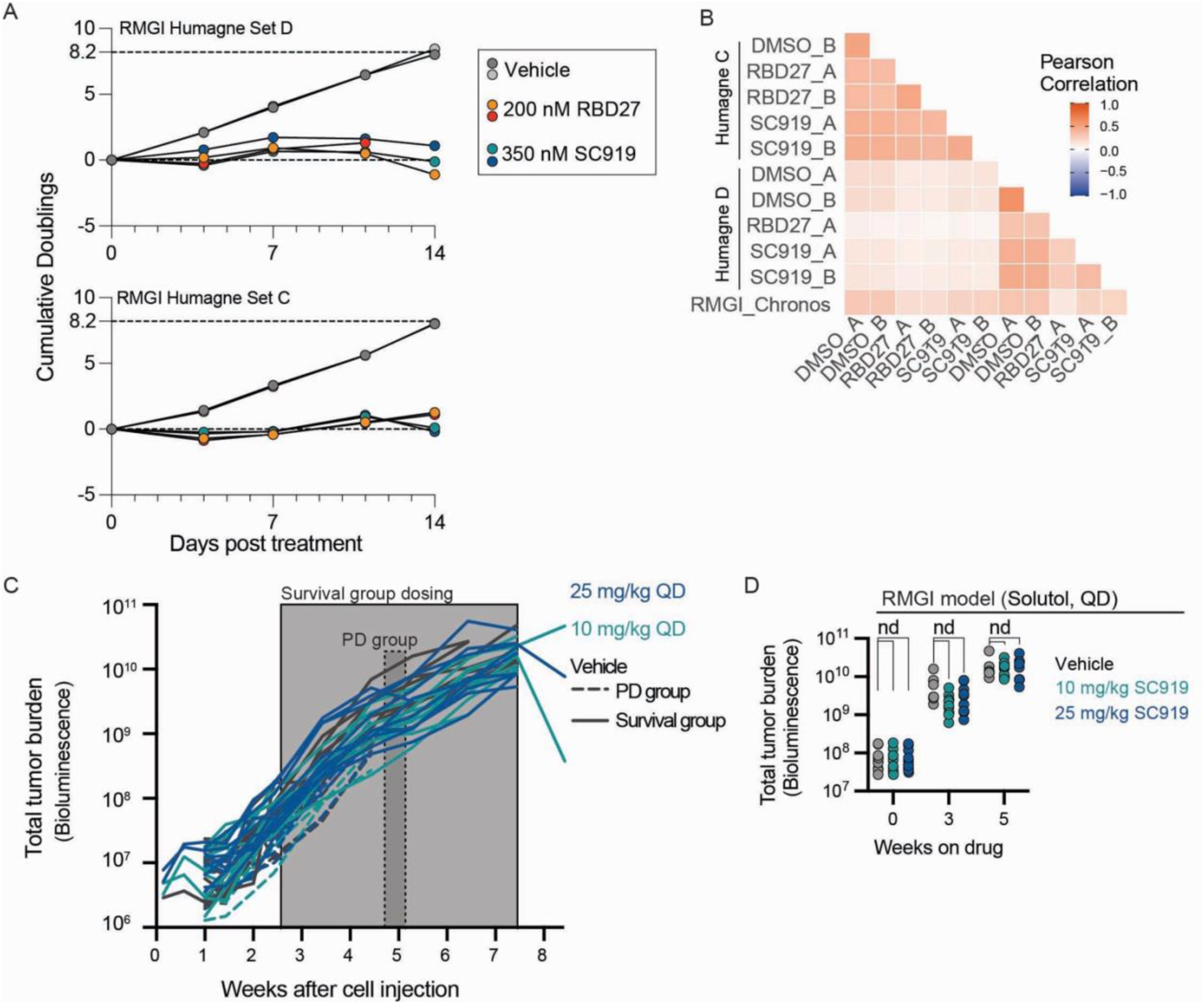
XPR1 and IP6K inhibitors share mechanisms of sensitivity but have unique mechanisms of resistance. A, Population growth throughout the genome scale modifier screen. The Humagne library is split into two sets (C and D), each containing a single, two-sgRNA array targeting each gene. After transduction at 1,000X coverage, selection, and expansion for several weeks, the cells were exposed to 200 nM RBD27 or 350 nM SC919 for two weeks. The cells were counted at the indicated timepoints, and cumulative population doublings were calculated. B, Comparison of replicate correlations in the genome-scale modifier screen. After sequencing the sgRNA barcodes from each replicate and deconvoluting the sequencing data into a read count matrix per condition and per sgRNA array, the correlation of gene essentiality and reads per million gene was compared. C, RMGI cells were injected IP into mice in a similar protocol to the OVISE study shown in Figure 3. After 20 days, animals were divided into survival and pharmacodynamic (PD) groups based on tumor burden. The survival groups were then randomized into vehicle and treatment groups and continuously dosed for 5 weeks using SC919 formulated in 10% DMSO and 10% Solutol at either 10 or 25 mg/kg. The PD group animals were treated on week five for 5 days prior to sacrifice. D, Summary of tumor burden and bioluminescence at various timepoints from panel C. A one-way ANOVA test revealed no significant differences between the untreated and treatment groups.

## REFERENCES

1. Bondeson, D. P. et al. Phosphate dysregulation via the XPR1–KIDINS220 protein complex is a therapeutic vulnerability in ovarian cancer. Nat. Cancer 3, 681–695 (2022).

2. Banerjee, S., Drapkin, R., Richardson, D. L. & Birrer, M. Targeting NaPi2b in ovarian cancer. Cancer Treat Rev 112, 102489 (2023).

3. Gerber, D. E. et al. Phase Ia Study of Anti-NaPi2b Antibody-Drug Conjugate Lifastuzumab Vedotin DNIB0600A in Patients with Non-Small Cell Lung Cancer and Platinum-Resistant Ovarian Cancer. Clin Cancer Res 26, 364–372 (2020).

4. Bodyak, N. D. et al. The Dolaflexin-based Antibody-Drug Conjugate XMT-1536 Targets the Solid Tumor Lineage Antigen SLC34A2/NaPi2b. Mol Cancer Ther 20, 896–905 (2021).

5. Forster, I. C., Hernando, N., Biber, J. & Murer, H. Phosphate transporters of the SLC20 and SLC34 families. Mol Asp. Med 34, 386–395 (2013).

6. Kim, S., Bhandari, R., Brearley, C. A. & Saiardi, A. The inositol phosphate signalling network in physiology and disease. Trends Biochem. Sci. 49, 969–985 (2024).

7. Zhou, W. et al. Kidney glycolysis serves as a mammalian phosphate sensor that maintains phosphate homeostasis. J Clin Invest 133, (2023).

8. Kritmetapak, K. & Kumar, R. Phosphate as a Signaling Molecule. Calcif Tissue Int 108, 16–31 (2021).

9. Li, X. et al. Control of XPR1-dependent cellular phosphate efflux by InsP8 is an exemplar for functionally-exclusive inositol pyrophosphate signaling. Proc Natl Acad Sci U A 117, 3568–3574 (2020).

10. Li, X. et al. Homeostatic coordination of cellular phosphate uptake and efflux requires an organelle-based receptor for the inositol pyrophosphate IP8. Cell Rep 43, 114316 (2024).

11. López-Sánchez, U. et al. Characterization of XPR1/SLC53A1 variants located outside of the SPX domain in patients with primary familial brain calcification. Sci Rep 9, 6776 (2019).

12. Wild, R. et al. Control of eukaryotic phosphate homeostasis by inositol polyphosphate sensor domains. Science 352, 986–990 (2016).

13. Wang, X. et al. KIDINS220 and InsP8 safeguard the stepwise regulation of phosphate exporter XPR1. Preprint at 10.1101/2025.01.17.633679 (2025).

14. Zhu, Q., Yaggi, M. F., Jork, N., Jessen, H. J. & Diver, M. M. Transport and InsP8 gating mechanisms of the human inorganic phosphate exporter XPR1. Nat. Commun. 16, 2770 (2025).

15. Bondeson, D. P. Insights into phosphate homeostasis regulation by XPR1. Nat. Struct. Mol. Biol. 1–3 (2024) doi:10.1038/s41594-024-01460-x.

16. Josifova, D. J. et al. Heterozygous KIDINS220/ARMS nonsense variants cause spastic paraplegia, intellectual disability, nystagmus, and obesity. Hum Mol Genet 25, 2158–2167 (2016).

17. Kwon, J. J. et al. Structure-function analysis of the SHOC2-MRAS-PP1C holophosphatase complex. Nature 609, 408–415 (2022).

18. Hanna, R. E., et al. Massively parallel assessment of human variants with base editor screens. Cell 184, 1064–1080.e20 (2021).

19. Lue, N. Z., et al. Base editor scanning charts the DNMT3A activity landscape. Nat. Chem. Biol. 19, 176–186 (2023).

20. Wang, W.-A. et al. Large-scale experimental assessment of variant effects on the structure and function of the citrate transporter SLC13A5. Sci. Adv. 11, eadx3011 (2025).

21. Marinko, J. T. et al. Folding and Misfolding of Human Membrane Proteins in Health and Disease: From Single Molecules to Cellular Proteostasis. Chem. Rev. 119, 5537–5606 (2019).

22. Legati, A. et al. Mutations in XPR1 cause primary familial brain calcification associated with altered phosphate export. Nat Genet 47, 579–581 (2015).

23. Zhang, J. et al. Gain-of-Function KIDINS220 Variants Disrupt Neuronal Development and Cause Cerebral Palsy. Mov. Disord. 39, 498–509 (2024).

24. Moritoh, Y. et al. The enzymatic activity of inositol hexakisphosphate kinase controls circulating phosphate in mammals. Nat Commun 12, 4847 (2021).

25. Battini, J. L., Rasko, J. E. & Miller, A. D. A human cell-surface receptor for xenotropic and polytropic murine leukemia viruses: possible role in G protein-coupled signal transduction. Proc Natl Acad Sci U A 96, 1385–1390 (1999).

26. Giovannini, D., Touhami, J., Charnet, P., Sitbon, M. & Battini, J.-L. Inorganic Phosphate Export by the Retrovirus Receptor XPR1 in Metazoans. Cell Rep 3, 1866–1873 (2013).

27. Yu, C. et al. High-throughput identification of genotype-specific cancer vulnerabilities in mixtures of barcoded tumor cell lines. Nat. Biotechnol. 34, 419–423 (2016).

28. Corsello, S. M. et al. Discovering the anticancer potential of non-oncology drugs by systematic viability profiling. Nat. Cancer 1, 235–248 (2020).

29. DeWeirdt, P. C. et al. Optimization of AsCas12a for combinatorial genetic screens in human cells. Nat Biotechnol 39, 94–104 (2021).

30. Carreras-Puigvert, J. et al. A comprehensive structural, biochemical and biological profiling of the human NUDIX hydrolase family. Nat. Commun. 8, 1541 (2017).

31. Kim, G. et al. Pools of independently cycling inositol phosphates revealed by pulse labeling with 18O-water. 2024.05.03.592351 Preprint at 10.1101/2024.05.03.592351 (2024).

32. Haykir, B. et al. The Ip6k1 and Ip6k2 Kinases Are Critical for Normal Renal Tubular Function. J Am Soc Nephrol 10.1681/ASN.0000000000000303 (2024) doi:10.1681/ASN.0000000000000303.

33. Ansermet, C. et al. Renal Fanconi Syndrome and Hypophosphatemic Rickets in the Absence of Xenotropic and Polytropic Retroviral Receptor in the Nephron. J Am Soc Nephrol 28, 1073–1078 (2017).

